# Blood mitochondrial health markers cf-mtDNA and GDF15 in human aging

**DOI:** 10.1101/2025.01.28.635306

**Authors:** Caroline Trumpff, Qiuhan Huang, Jeremy Michelson, Cynthia C. Liu, David Shire, Christian G. Habeck, Yaakov Stern, Martin Picard

**Author notes:** Corresponding authors. Department of Psychiatry, Division of Behavioral Medicine, Columbia University Medical Center, New York, USA.

## Abstract

Altered mitochondria biology can accelerate biological aging, but scalable biomarkers of mitochondrial health for population studies are lacking. We examined two potential candidates: 1) *cell-free mitochondrial DNA* (cf-mtDNA), a marker of mitochondrial signaling elevated with disease states accessible as distinct biological entities from plasma or serum; and 2) *growth differentiation factor 15* (GDF15), an established biomarker of biological aging downstream of mitochondrial energy transformation defects and stress signaling. In a cohort of 430 participants aged 24-84 (54.2% women), we measured plasma and serum cf-mtDNA, and plasma GDF15 levels at two timepoints 5 years apart, then assessed their associations with age, BMI, diabetes, sex, health-related behaviors, and psychosocial factors. As expected, GDF15 showed a positive, exponential association with age (r=0.66, p<0.0001) and increased by 33% over five years. cf-mtDNA was not correlated with GDF15 or age. BMI and sex were also not related to cf-mtDNA nor GDF15. Type 2 diabetes was only positively associated with GDF15. Exploring potential drivers of systemic mitochondrial stress signaling, we report a novel association linking higher education to lower age-adjusted GDF15 (r=-0.14, p<0.0034), both at baseline and the 5-year follow up, highlighting a potential influence of psychosocial factors on mitochondrial health. Overall, our findings among adults spanning six decades of lifespan establish associations between age, diabetes and GDF15, an emerging marker of mitochondrial stress signaling. Further studies are needed to determine if the associations of blood GDF15 with age and metabolic stress can be moderated by psychosocial factors or health-related behaviors.

## 1. Introduction

Individuals differ in their pace of biological aging and health span ^1^, but the underlying molecular mechanisms explaining these differences remain unclear. Accelerated aging might arise from altered mitochondrial biology^2,3^, which could stem from behavioral changes ^4,5^, but scalable, non-invasive biomarkers of mitochondrial health to address this question in population studies are lacking. Two potential candidate mitochondrial health biomarkers are circulating cell-free mitochondrial DNA (cf-mtDNA) and growth differentiation factor 15 (GDF15).

Mitochondria are the only mammalian organelle besides the nucleus to contain their own genome, populating every nucleated cell in the body. Each mitochondrion contains multiple copies of the 16.6kb-long circular mitochondrial DNA (mtDNA) bound in protein-DNA complexes typically contained within mitochondria ^6,7^. But the mtDNA is also detectable extracellularly (i.e., outside the cell) in most body fluids, including blood. Previous studies reported that blood cf-mtDNA levels are elevated in aging ^8^ and in diseases states including trauma ^9^, inflammatory diseases ^10–14^, diabetes ^15–18^, as well as cancers, myocardial infarction, and sepsis ^19–23^. Plasma and serum cf-mtDNA represent distinct biological entities: plasma is closest to what circulates in the bloodstream at the time of draw, whereas serum includes mtDNA and related mtDNA-containing particles released during coagulation ^24^. As a result, these types of cf-mtDNA are akin to different types of circulating cholesterol (e.g., LDL, HDL) ^25^ and in preliminary studies are differently associated with health-related parameters ^26^. This calls for larger studies to understand the associations of different forms of cf-mtDNA, including plasma or serum-derived cf-mtDNA, with aging and disease.

In the aging body, the brain must be aware of systemic energetic stress ^5^. Tissue metabolic stress is conveyed to the brain by the circulating metabokine-cytokine GDF15 ^27^. GDF15 is the leading candidate biomarker for primary genetic mitochondrial diseases ^28–33^, and it is also the most robustly elevated circulating protein with human aging ^34,35^ – possibly pointing to shared underlying biology. GDF15 is also elevated in several chronic physical illnesses including diabetes ^36^, cardiovascular diseases ^37^, cancer ^38^, Alzheimer’s disease ^39,40^ autoimmune diseases ^41^. In studies of multimorbidity in thousands of individuals, GDF15 emerges as the top discriminatory plasma protein associated with multiple current ^42^ and future clinical illnesses ^40^. It also is among the best prognostic mortality indicators in clinical populations and older adults ^34,43^. Psychosocial stress might also affect GDF15, a recent study found that poverty influence GDF15 levels and interact with mortality risk ^44^. Taken together, the rapidly growing body of literature suggests that GDF15 may play a role in systemic energy homeostasis, resilience, and health and represent a potential biomarker of the stress-accelerated aging cascade.

While GDF15 has been shown to increase within person over 5-10 year periods ^45^, no prior studies have evaluated whether plasma or serum forms of cf-mtDNA change over time. Here we assayed plasma and serum cf-mtDNA, in parallel with GDF15, from the Reference Ability Neural Network/Cognitive Reserve (RANN/CR) study of adults spanning 6 decades of the adult lifespan. ^46^ We report associations between blood cf-mtDNA (plasma and serum) and GDF15 levels in relation to age, sex, BMI, diabetes and psychosocial and lifestyle factors.

## 2. Results

### 2.1. Plasma vs serum forms of cf-DNA

In total, we assayed 1003 samples from 430 individuals, including 430 plasma samples at the baseline timepoint, and 297 plasma and 276 serum samples 5 years later. *Plasma* was available both at baseline and 5-year follow up, whereas *serum* was collected only at 5-year follow up. The demographic information and participants’ characteristics are shown in Table 1.

**Table 1.** Demographics and Participant Characteristics Table.

|  | **Baseline** | **Follow-up** |
| --- | --- | --- |
| Number of Participants | 430 | 298 |
| Sex, No. (%) |  |  |
| Female | 223 (51.9) | 156 (52.3) |
| Male | 185 (43.0) | 138 (46.3) |
| Age (years) * | 55.0 (16.6) | 58.9(16.3) |
| Race<sup>†</sup>, No. (%) |  |  |
| Asian | 12 (2.8) | 8 (2.7) |
| Black or African American | 110 (25.6) | 68 (22.8) |
| Native Hawaiian or Other Pacific Islander | 5 (1.2) | 4 (1.3) |
| White | 268 (62.3) | 198 (66.4) |
| Other | 24 (5.6) | 14 (4.7) |
| Body Mass Index * | 26.5 (4.9) | 25.7 (4.7) |
| Yearly household income, No. (%) <sup>‡</sup> |  |  |
| < \$5,000 | 9 (2.1) | 3 (1.0) |
| \$5,000 – \$11,999 | 18 (4.2) | 8 (2.7) |
| \$12,000 – \$15,999 | 10 (4.3) | 8 (2.7) |
| \$16,000 – \$24,999 | 18 (4.2) | 6 (2.0) |
| \$25,000 – \$34,999 | 22 (5.1) | 14 (4.7) |
| \$35,000 – \$49,999 | 30 (7.0) | 28 (9.4) |
| \$50,000 – \$74,999 | 61 (61) | 40 (13.4) |
| \$75,000 – \$99,999 | 34 (7.9) | 33 (11.1) |
| > \$100,000 | 96 (22.3) | 88 (29.5) |
| Reported “don’t know” | 17 (4.0) | 10 (3.4) |
| Education (years) * | 15.9 (2.4) | 16.0 (2.3) |
| Functional Activity * | 0.37 (0.9) | 1.26 (1.5) |
| Mild Physical Activity (METs) * | 7.7 (8.8) | 9.8 (44.6) |
| Moderate Physical Activity (METs) * | 12.8 (10.5) | 13.8 (12.0) |
| Strenuous Physical Activity (METs) * | 8.5 (9.7) | 16.7 (137.8) |
| Total Physical Activity (METs) * | 27.8 (20.3) | 40.6 (152.9) |
| Social Support * | 2.7 (0.5) | 2.6 (0.6) |
| Social Network Size * | 10.5 (9.1) | 10.6 (10.7) |
| Total Leisure activity * | 35.6 (4.4) | 35.5 (5.3) |
| Leisure Activity Change (6 months) * | 2.5 (0.6) | 2.4 (0.8) |
| Leisure Activity Change (10 years) * | 2.3 (0.8) | 2.4 (0.8) |
| Perceived Stress Scale (PSS) * | NA | 11.8 (6.4) |

\* Data shown as mean (standard deviation)
† Not all participant reported race
‡ Not all participant reported income

Plasma and serum contains distinct forms of cf-mtDNA ^47^ that correlate differently with health-related parameters ^26^. This motivated us to directly compare plasma and serum cf-mtDNA in the same individuals, over time. Cf-mtDNA can be released in the circulation via both active release mechanisms (e.g. vesicles) and possibly cell death, while the cell-free nuclear DNA (cf-nDNA) is likely only released after cell death ^47–51^. This makes the mitochondrial:nuclear ratio (cf-mtDNA/cf-nDNA) potentially meaningful to isolate the contribution of mitochondrial release.

In line with previous reports, cf-mtDNA levels were on average 5.1-fold higher (p<0.0001, Fig. S1A) in serum vs plasma. Similarly, cf-nDNA levels were on average 20.3-fold higher in serum vs plasma (p<0.0001, Fig S1B). Serum and plasma cf-mtDNA showed a weak positive correlation, confirming the distinct biological nature of these analytes (Fig. S1D). Serum and plasma cf-nDNA were not correlated (Fig. S1E). cf-mtDNA/nDNA ratio, were on average 1.7-fold higher in plasma vs serum (p<0.0001, Fig. S1C), in line with the idea that serum contains higher levels of mtDNA derived from cell death than plasma. There was a weak negative correlation between the serum and plasma cf-mtDNA/nDNA ratio (Fig. S1F). Different proportions of cf-mtDNA may be retained during the serum or plasma coagulation process, potentially accounting for the observed discrepancy between the two sample types ^26^.

Comparing mitochondrial and nuclear signals, at baseline there was a weak negative association between cf-mtDNA and cf-nDNA in plasma (r=-0.11, p=0.025, Fig. S1G) while moderate positive associations were found in both plasma and serum at the 5-year follow-up visit (r=0.35, p<0.0001 and r=0.51, p<0.0001 respectively, Fig. S1H-I). Thus, overall, these results are consistent with a distinct regulation of cf-mtDNA and cf-nDNA, but also suggest that individuals with higher cf-mtDNA also tend to have high cf-nDNA.

### 2.2 GDF15 and cf-mtDNA

Next, we evaluated the associations between cf-mtDNA and GDF15. No associations were found between plasma cf-mtDNA and GDF15 levels at baseline (Fig. S1J). There was a weak positive association between cf-mtDNA and GDF15 at the follow-up visit with plasma (r=0.15, p=0.0083, Fig. S1K) but not with serum cf-mtDNA (Fig. S1L).

### 2.3. cf-DNA by age and sex

Cross-sectionally, plasma cf-mtDNA was not consistently associated with age. A weak negative association was found at baseline (r=-0.13, p=0.007, Fig. 1B) but not at the follow-up (r=0.11, p=0.052, Fig. S2A). However, plasma cf-mtDNA levels were found to strongly increase by an average of 3.7-fold over 5 years (p<0.0001, Fig. 1C and Fig S2B). Regarding the within-person stability, cf-mtDNA levels were only weakly correlated between baseline and follow-up visits (r=0.15, p=0.037, Fig. 1D), consistent with the dynamic inducibility of plasma cf-mtDNA ^52,53^. Individuals with higher plasma cf-mtDNA levels at baseline tended to show lower change from baseline to follow-up (r=-0.15, p=0.04, Fig. 1E).

**Figure 1:**
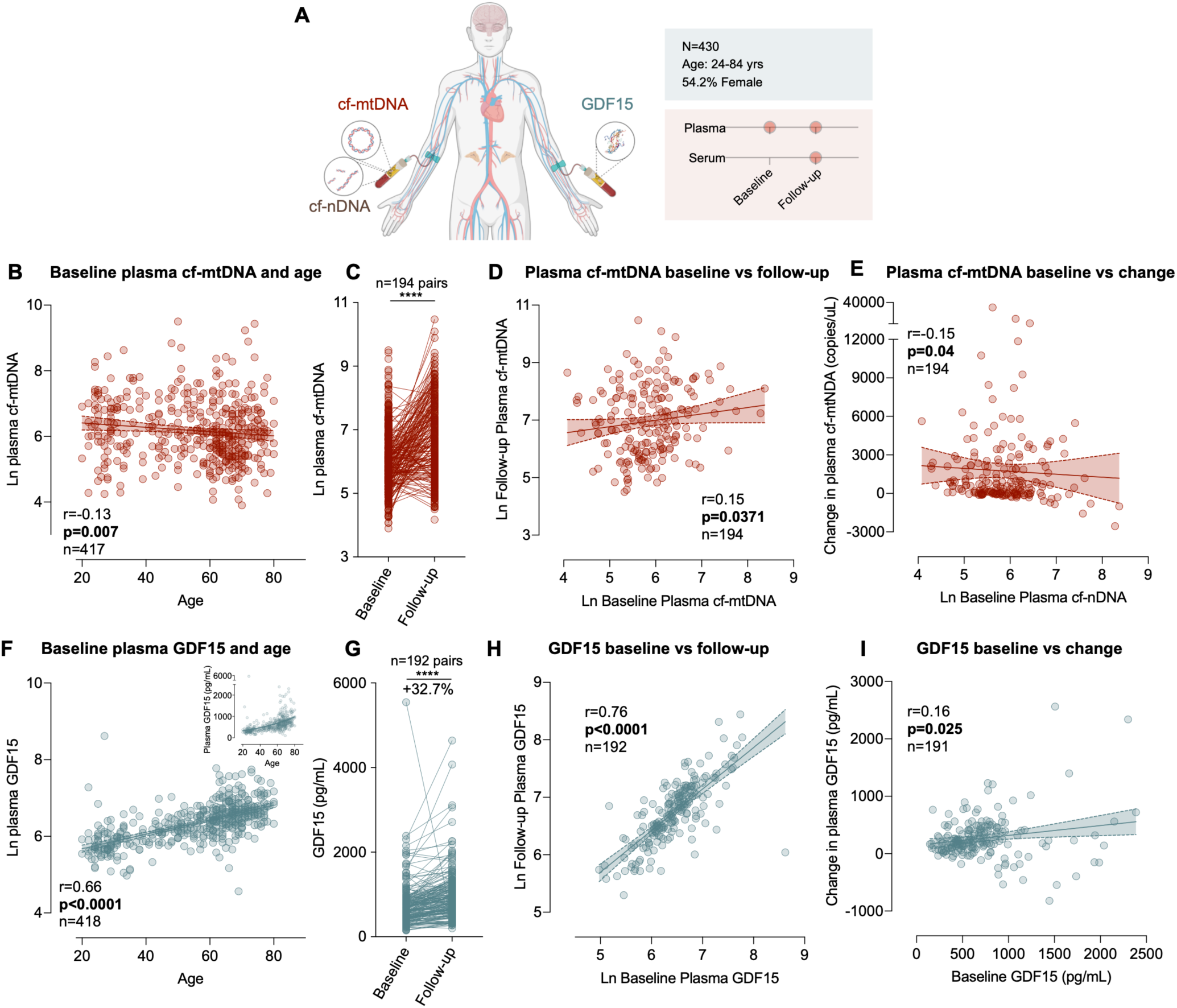
Plasma cf-mtDNA and GDF15 association with age and within-individual trajectories over 5 years. **(A)** Study design: Plasma and serum samples from the RANN-CR study were used to measure cf-mtDNA and GDF15 levels at baseline and 5-year follow-up. **(B)** Scatterplot of the association between plasma cf-mtDNA levels and age at baseline. For the correlation with age in the follow-up visit see Figure S2A.(C) Change in cf-mtDNA levels from baseline to follow-up, for raw concentrations in copies/ul see Fig. S2B. **(D)** Scatterplot of the association between plasma cf-mtDNA levels at baseline and follow-up. **(E)** Scatterplot of the association between baseline and change in plasma cf-mtDNA levels from baseline to follow-up.(F, **G, H,** I) Same as in **B,C,D** but for plasma GDF15. Insert graph show baseline GDF15 concentration in pg/ml demonstrating the exponential nature of the association with age. For the correlation between plasma GDF15 and age in the follow-up visit see Figure S2F. P-value and effect sizes from **(B,D,E,F,H,I)** Spearman rho correlation and **(C,G)** Wilcoxon paired t-test. *p<0.05, **p<0.01, ***p<0.001, ****p<0.0001.

Plasma cf-nDNA levels, on the other hand, showed a weak positive cross-sectional association with age at baseline (r=0.18, p=0.0002, Fig. S2D), although not at the follow-up visit (r=0.011, p=0.86, Fig. S2E). Plasma cf-nDNA levels were moderately correlated within individuals between baseline and follow-up (r=0.44, p<0.0001, Fig. S2F), demonstrating a greater stability for cf-nDNA than cf-mtDNA. In contrast to the remarkable 5-year elevation in cf-mtDNA, there was no significant change in plasma cf-nDNA levels over the 5-year period (Fig. S2G), confirming a selective change in plasma cf-mtDNA over time (Fig. 1C).

Plasma cf-mtDNA/cf-nDNA ratios presented contrasting association with age, showing a negative association at baseline (r=-0.20, p<0.0001, Fig.S2H) and a positive association at follow-up (r=0.15, p=0.011, Fig. S2I). Plasma cf-mtDNA/cf-nDNA ratios were not significantly associated between baseline and follow-up (r=-0.13, p=0.073, Fig. S2J). Similarly to cf-mtDNA, the cf-mtDNA/cf-nDNA ratio increased 3.1-fold over the 5-year period (p<0.0001, Fig. S2K), suggesting that the observed cf-mtDNA increase was associated to active mtDNA release mechanisms, rather than general cell damage or turnover. Individuals with higher plasma cf-mtDNA/cf-nDNA levels at baseline showed higher change from baseline to follow-up (r=0.45, p<0.0001, Fig. S2L).

In serum, no correlation was found between age and serum cf-mtDNA, cf-nDNA, nor cf-mtDNA/cf-nDNA ratio at the follow-up visit (Fig. S3). Males and females exhibited similar plasma or serum cf-mtDNA, cf-nDNA, and cf-mtDNA/cf-nDNA ratio levels (Fig. S4A-C, S4E-G), ruling out a sex difference in this cohort.

### 2.4. GDF15 levels by age and sex

As expected, cross-sectionally, plasma GDF15 levels were positively associated with age at both baseline (r=0.66, p<0.0001, Fig.1F) and 5 years later (r=0.75, p<0.0001, Fig. S2C). These large effect size associations are exponential, such that GDF15 increases more rapidly as individuals get older. Over the 5-year follow up, GDF15 increased by an average of 32.7% (p<0.0001, Fig. 1G). Baseline and follow-up GDF15 levels were strongly correlated within individuals (r=0.76, p<0.0001, Fig. 1H), reflecting the trait nature of this biomarker. Higher baseline GDF15 level also predicted larger change in GDF15 from baseline to follow-up (r=0.16, p=0.025, Fig. 1I). Females and males showed similar GDF15 levels both at the baseline and follow-up visits (Fig. S4D and S4H).

### 2.5. cf-DNA levels, GDF15 levels and metabolic health

In this dataset, information on BMI and type two diabetes mellitus diagnosis was collected at both baseline and follow-up visits and was used here as a proxy for assessing metabolic health. However, the sample size for individual with diabetes is small, which may lead to an overestimation of the effect size.

Plasma cf-mtDNA showed no consistent associations with BMI (Fig. 2A) or diabetes (Fig. 2B) at baseline and follow-up. Similarly, plasma cf-nDNA levels and cf-mtDNA/cf-nDNA ratios were not associated with neither BMI nor diabetes (Fig. S5A-B, Fig. S5C-S5D). In serum, no associations with diabetes were found in cf-mtDNA, cf-nDNA, or cf-mtDNA/cf-nDNA ratio levels (Fig. S5F, H, J). In relation to BMI, no association was found in serum cf-mtDNA levels (Fig.S5E). Interestingly, a weak positive correlation was found in serum cf-nDNA levels (r=0.20, p=0.007, Fig. S5G) indicating that nuclear DNA levels may be elevated in individuals with higher BMI, potentially reflecting greater cell turnover or tissue damage associated with adiposity. The weak negative correlation between the serum cf-mtDNA/cf-nDNA ratio and BMI (r=-0.23, p=0.0016, Fig. S5I) further supports that in individuals with higher BMI, the relative presence of cf-mtDNA compared to cf-nDNA is reduced.

**Figure 2:**
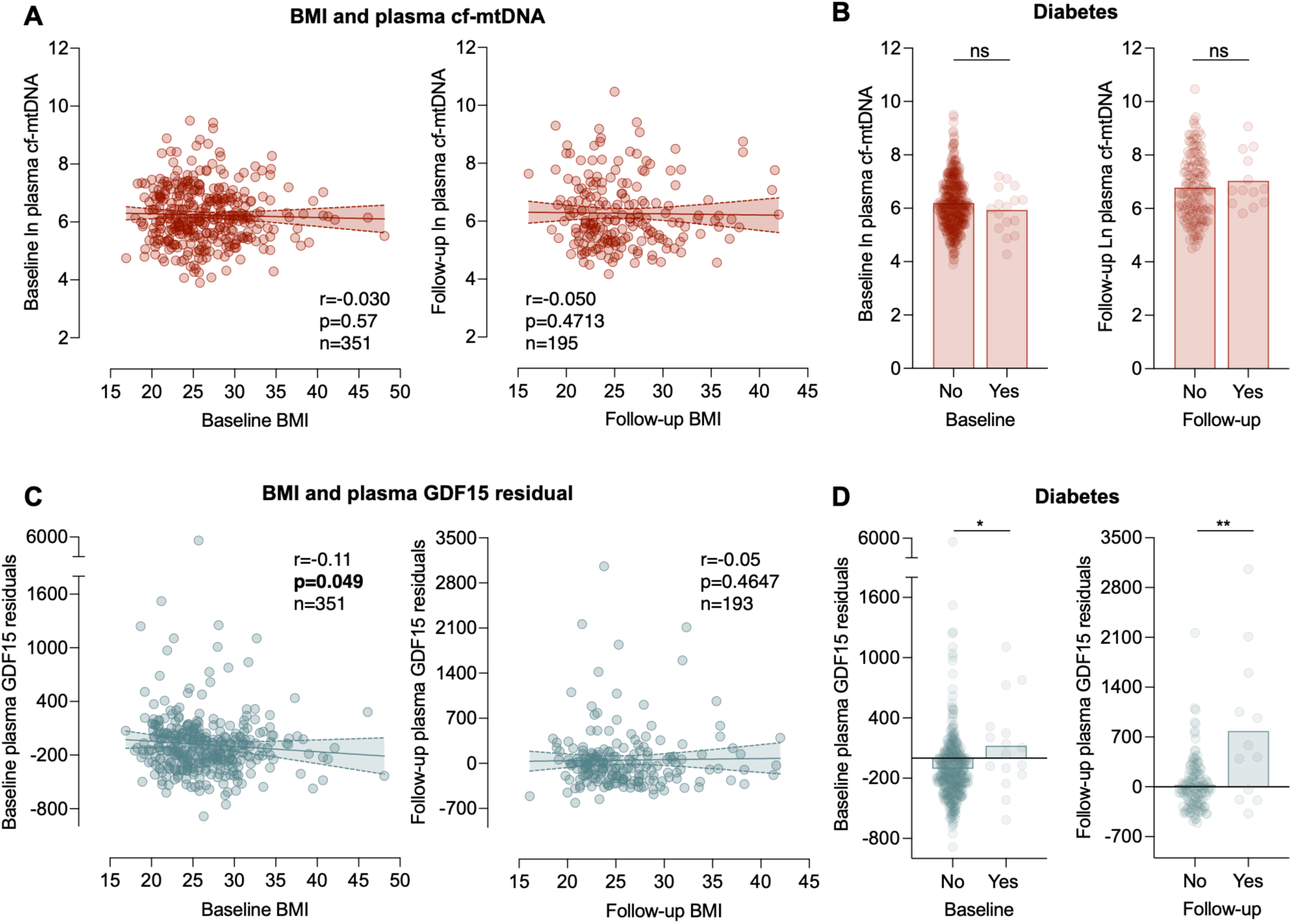
GDF15 and cf-mtDNA association with BMI and diabetes. **(A)** Scatterplot of the association between participant’s plasma cf-mtDNA levels and BMI at the baseline (left) and follow-up (right) visit. **(B)** Difference in plasma cf-mtDNA levels by diabetic status at the baseline (left, n=401 vs 16) and follow-up (n=135 vs 13) visit. **(C)** Same as in A but for plasma GDF15 at baseline (left) and follow-up(right) visit. **(D)** Same as in B but for plasma GDF15 at baseline (left, n=402 vs 16) and follow-up (right, n=134 vs 12) visit. P-value and effect sizes from **(A,C)** Spearman rho correlation and **(B,D)** Mann-Whitney t-test. *p<0.05, **p<0.01, ***p<0.001, ****p<0.0001.

Regarding GDF15, at baseline, BMI showed a weak negative association with age-adjusted plasma GDF15 levels (r=-0.11, p=0.049, Fig.2C (left)). At the 5-year follow-up, where the sample size was smaller, we found no significant association with BMI. Both at baseline and at follow-up, individuals diagnosed with type 2 diabetes exhibited significantly higher GDF15 compared to those without a diagnosis (p=0.011, Fig. 2D left, p=0.007, Fig. 2D right). However, again, this effect size may be overestimated given the small number of individuals (n=16 at baseline, n=12 at follow-up) with diabetes in this cohort.

### 2.6. 5-year change in cf-DNA and GDF15 levels with age and sex

Since plasma cf-mtDNA, cf-mtDNA/cf-nDNA ratio, and age-adjusted GDF15 levels increased over time, we investigated participant characteristics that explained inter-individual difference in the magnitude of this increase. The magnitude of 5-year increase in plasma cf-mtDNA levels and cf-mtDNA/cf-nDNA ratio was not associated with sex or age at baseline (Fig. S6A-F). We did not find a sex difference (Fig. S6G) in the magnitude of change in age-adjusted GDF15 levels. The magnitude of increase in age-adjusted GDF15 was positively associated with age (r=0.40, p<0.0001, Fig. S6H) indicating that older individuals have a larger increase in GDF15 levels with time.

### 2.7. Associations between 5-year change in plasma cf-DNA and GDF15 levels and metabolic health

Next, we explored the relationship between BMI and type 2 diabetes diagnosis, and the magnitude of the 5-year change in plasma cf-DNA and GDF15 levels. For BMI, we did not find any association between change in plasma cf-mtDNA or cf-mtDNA/cf-nDNA ratio and BMI at baseline or follow-up (Fig.S7A, S7E). In contrast, while no association was observed between changes in GDF15 levels and BMI at follow-up (Fig. S7J). higher baseline BMI was associated with a greater increase in age-adjusted GDF15 levels from baseline to follow-up (r=0.20, p=0.013, Fig. S7I).

Baseline levels of plasma cf-mtDNA, cf-mtDNA/cf-nDNA ratio or age-adjusted GDF15 levels were not associated with diabetes status at follow-up (Fig. S7B, S7F, S7J). Similarly, the magnitude of change in plasma cf-mtDNA levels or cf-mtDNA/cf-nDNA ratios was not related to the diabetic status at baseline or follow-up (Fig. S7C, S7G, Fig. S7D, S7H). In contrast, compared to those without a diagnosis, we found that individuals diagnosed with diabetes at follow-up had a larger increase in age-adjusted GDF15 levels from baseline to follow-up (p=0.028, Fig. S7K), suggesting that GDF15 changes may reflect metabolic stress underlying diabetes risk. Additionally, individuals diagnosed with diabetes at baseline also exhibited a greater change in age-adjusted GDF15 levels (p=0.0001, Fig. S7L), linking systemic glucose intolerance and insulin resistance with energetic and mitochondrial stress signaling ^54^.

### 2.8. Associations between cf-mtDNA, GDF15 and behavioral and psychosocial factors

Finally, we explored the associations between behavioral and psychosocial factors and cf-mtDNA and GDF15 levels. Given emerging literature on the interconnection of psychosocial factors, the biology of energy and mitochondria ^55^, we conducted these analyses for hypothesis generation, exploring associations de novo without prior hypotheses, and thus used uncorrected p-values.

At baseline (Fig. 3A), individuals who perform more mild physical activity (r=-0.14, p=0.0044, Fig. 3A’) and have participated in more years of education (r=-0.097, p=0.047, Fig.3A”) had lower plasma cf-mtDNA levels. Similarly, the plasma cf-mtDNA/cf-nDNA ratio at baseline (Fig. 3B) were negatively associated with mild physical activity (r=-0.13, p=0.021, Fig. 3B’), adding specificity to this finding. At the follow-up visit, no significant associations were observed between plasma cf-mtDNA levels or the cf-mtDNA/cf-nDNA ratio, and behavioral or psychosocial factors (Fig. S7A-B). At the follow-up visit, serum cf-mtDNA levels (Fig. S8C) showed a negative association with moderate physical activity (r=-0.14, p=0.039, Fig. S8C’). No significant associations were observed between the serum cf-mtDNA/cf-nDNA ratio and behavioral or psychosocial factors (Fig. S8D).

**Figure 3:**
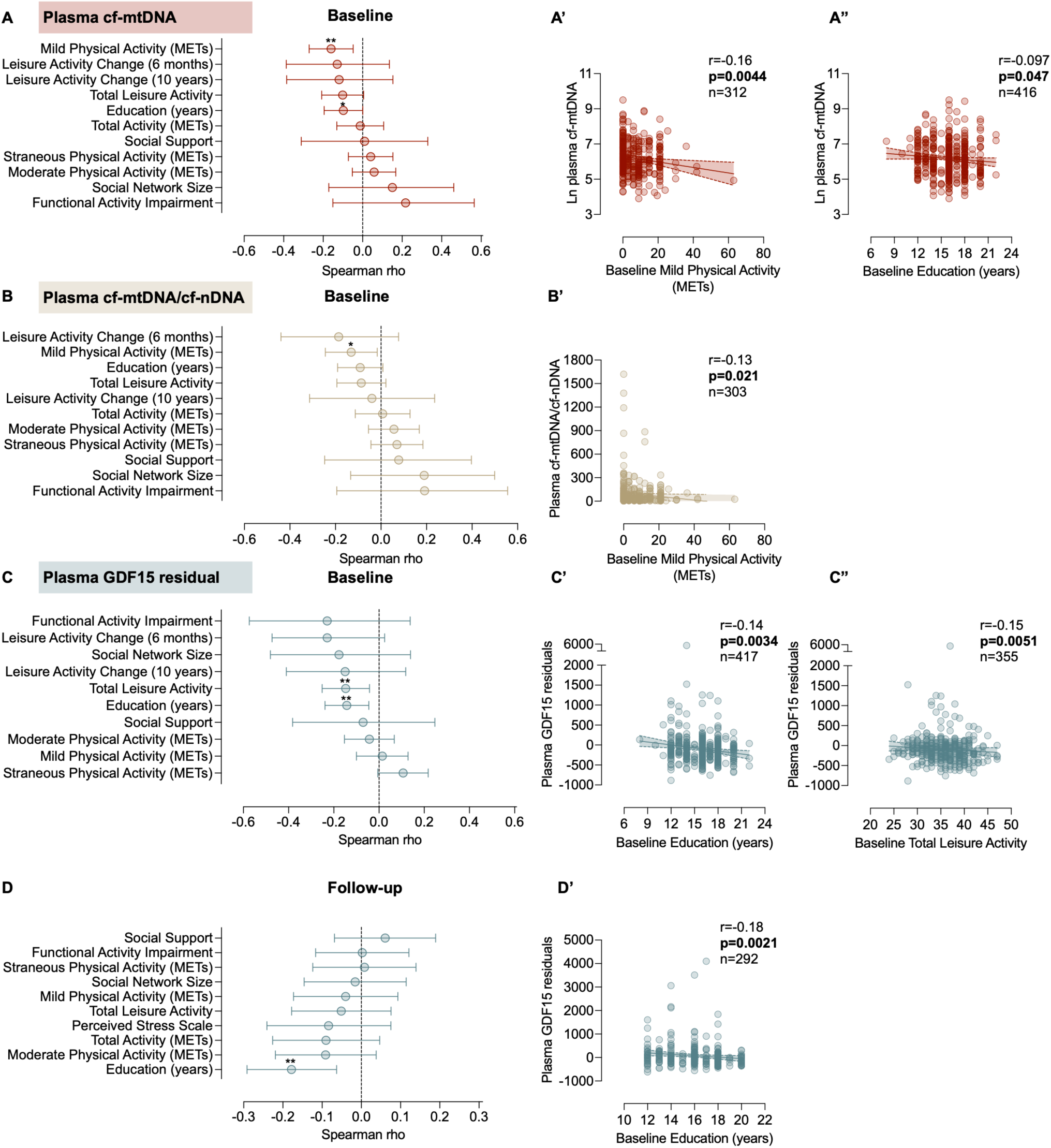
Associations between plasma GDF15 levels and behavioral and psychosocial factors. **(A)** Associations between baseline plasma cf-mtDNA levels with behavioral and psychosocial factors. See **(A’)** and **(A“)** for statistically significant correlations with Mild Physical Activity (METs) and Education respectively. **(B)** Associations between baseline plasma cf-mtDNA/cf-nDNA ratio with behavioral and psychosocial factors. See **(B’)** for statistically significant correlations with Mild Physical Activity (METs). **(C)** Associations between baseline plasma GDF15 levels with behavioral and psychosocial factors. See **(C’)** and **(C”)** for statistically significant correlations with Education and Total Leisure Activity respectively. **(D)** Same as **C** but for plasma GDF15 levels at follow-up. See **(D’)** for statistically significant correlation with Education. P-value and effect sizes from Spearman rho correlation. See Supplemental Table 1 for sample size of each measurement. *p<0.05, **p<0.01, ***p<0.001, ****p<0.0001.

Regarding GDF15 (Fig. 3C-D), age-adjusted plasma GDF15 levels was negatively associated with total leisure activity (r=-0.15, p=0.0051, Fig. 3C”) at baseline, but not at follow-up. Higher level of education was found to be negatively associated with age-adjusted plasma GDF15 levels both at baseline (r =-0.14, p=0.0034, Fig. 3C’) and follow-up (r=-0.18, p=0.0021, Fig.3D’), suggesting that higher education levels might be confer protection against altered mitochondrial biology or other forms of energetic stress upstream of GDF15 release.

Finally, behavioral and psychosocial factors were not related to 5-year changes in neither cf-mtDNA (Fig. S8E) nor GDF15 (Fig. S8F).

## 3. Discussion

This study demonstrates that two mitochondrial signaling markers — circulating plasma cf-mtDNA and GDF15 — increase over a 5-year period. In relation to the biology of cell-free DNA, our study extends prior small studies ^56,57^ into a considerably larger sample, contrasting plasma and serum biofluids, and both mitochondrial and nuclear DNA. Combined with the robust age-related elevation in GDF15, these findings expand our understanding of how these markers are (un)related, depending on the measurement approach, representing a step towards developing useful, minimally invasive markers to assess mitochondrial health and signaling in aging populations.

The coagulation-independent plasma cf-mtDNA was not associated with circulating nuclear DNA; but people with higher levels of serum cf-mtDNA (measured after blood coagulation) had more cf-nDNA. In line with this concept and with prior studies ^56,57^, cf-mtDNA levels were higher in serum than plasma, but the cf-mtDNA/cf-nDNA ratio was larger in plasma than serum. This supports the idea that serum cf-mtDNA may partly come from cell breaking during coagulation, while plasma cf-mtDNA may more unambiguously represent cell-free mtDNA found in circulation ^47,58^. As previously reported by our group ^26^, we did not find evidence of a significant positive association between plasma and serum cf-mtDNA, further suggesting they biological forms of cell-free mitochondrial components represent distinct populations, or biological subsets of cf-mtDNA.

Regarding the relationship between cf-mtDNA and GDF15, in line with prior reports^59^, we found no robust nor consistent associations between the two biomarkers. Although both markers are understood to be “stress inducible” ^52,53,60^. This result suggests they may be independently regulated and reflect distinct cellular stress responses.

How cf-mtDNA levels change with aging has not been studied in detail. Only one prior report studied cf-mtDNA-aging relationship across separate populations of adults spanning ages 21 to ≥90 years old (N=516). They reported a cross-sectional positive association between plasma cf-mtDNA levels and age^8^. The current study did not replicate this finding cross-sectionally. However, our observed 5-year 3.7-fold increase in plasma cf-mtDNA is consistent with this earlier finding. While the biological specificity of the plasma cf-mtDNA increase was indicated by the lack of increase of plasma cf-nDNA, the lack of cross-sectional association with age in this study suggests that these results mtDNA should be considered with caution and need to be replicated in future studies. Future studies should also assess the association between age and cf-mtDNA considering the distinct biological forms of cf-mtDNA. Contrasting with our findings, a study showed a decrease in mtDNA content of extracellular vesicle isolated from plasma with age ^61^, further reinforcing the notion that similar to the LDL/HDL forms of circulating cholesterol, different cf-mtDNA forms (within vesicles, within mitochondria, or free-floating) may reflect distinct underlying biology.

In line with previous studies finding robust positive correlations between GDF15 and age ^35,45,62,63^, we also found a strong cross-sectional association between age and GDF15 levels. Consistent with a previous study ^45^, individuals with higher initial plasma GDF15 levels exhibited a greater increase in GDF15 over time. Interestingly, a larger change in GDF15 levels over time have been linked to increased mortality risk ^64^, suggesting that elevated GDF15 levels may serve as a marker of ongoing systemic stress, potentially reflecting the physiological decline commonly associated with aging.

We did not find evidence of a sex difference in baseline plasma and serum cf-mtDNA levels. This is similar to some previous reports ^65–67^ but also contrasts with other prior studies showing higher levels in men compared to women ^68,69^. We previously found either none or different direction of sex difference according to the biofluid analyzed (no difference with EDTA plasma or serum red top tube, higher in men in serum gold tube, higher in women in citrate plasma) ^26^. Regarding sex difference in GDF15, while previous report have found higher GDF15 levels in men ^70,71^, we did not find evidence of a sex difference in our cohort. However, on average, the 5-year change in GDF15 was larger in men than women, which is in line with the idea that men might age faster than women ^72^.

Regarding metabolic health, we previously found no association between BMI and plasma cf-mtDNA and a negative association when serum cf-mtDNA was measured from red top but not gold collection tubes ^26^. Here, we did not find evidence of an association neither for serum nor plasma cf-mtDNA and BMI. Contrasting with several previous studies showing elevated blood cf-mtDNA in diabetes ^15–18^, there were no significant elevation in plasma or serum cf-mtDNA levels in participants with type 2 diabetes. These results should be interpreted with caution due to the small sample size of participants diagnosed with type 2 diabetes in this study.

Prior reports found obesity to be associated with elevated GDF15 ^73–75^. In contrast, a study of genetically identical twins found that the twin with higher GDF15 levels presented a lower BMI^76^. A lower BMI was also associated with higher GDF15 levels in pregnant women^77^. Moreover, high GDF15 is linked to weight loss and cachexia in cancer, and neutralizing circulating GDF15 with a monoclonal antibody reduced the loss of muscle mass ^78^, suggesting that GDF15 contributes to weight loss. The present study of individuals across a wide age range found a weak negative association between age-adjusted GDF15 and BMI at baseline. However, this association was not observed at follow-up. This finding might be due to the smaller sample size at follow-up or might suggest that the relationship with BMI may not be consistent with age. We also found that people with greater BMI at baseline tend to have greater increase in GDF15 levels from baseline to follow-up, suggesting that BMI could influence the trajectory of GDF15 levels, possibly reflecting metabolic stress-related processes. And as in prior reports demonstrating elevated GDF15 in individuals with diabetes ^36,71^, we found elevated age-corrected GDF15 in study participants with type 2 diabetes at both timepoints, consistent with the notion that systemic insulin resistance and the resulting metabolic stress, which can compromise mitochondrial quality control^54^, is associated with elevated GDF15. Both the presence of type 2 diabetes at baseline and follow-up were associated with a larger 5 year increase in GDF15 levels, suggesting GDF15 may reflect metabolic stress underlying diabetes risk.

Because mitochondria produce the energy and signals necessary for stress adaptation, there is a direct connection between stress responses and mitochondrial biology ^79,80^. Several studies have found that psychological stress alter several aspects of mitochondria biology ^81–86^. Emerging evidence also suggest that greater positive psychosocial experience such as positive mood ^87^ or greater well-being ^88^ are associated with better mitochondrial health in immune and brain tissues, suggesting a connection between psychosocial factors and mitochondrial health. Here, we preliminarily explored this question by linking behavioral and psychosocial factors to cf-mtDNA and GDF15. In relation to cf-mtDNA, our most significant finding is that mild physical activity and higher education levels were associated with lower *plasma* cf-mtDNA (only at baseline, not at follow-up). These associations call for further studies. Regarding GDF15, the most significant finding was that higher level of education was associated with lower GDF15 levels at both study timepoints, in line with a previous study showing that poverty influences GDF15 levels ^44^. Using the human proteome atlas from >52,000 adults in the UK Biobank investigating the plasma proteome in health and disease ^89^, we also identified negative associations between plasma GDF15 levels and years of education (β=-0.28, CI=-0.41 to-0.15, SE = 0.068, p=3.98×10^-5^). Similarly, leisure-related activities, such as going to sports club and gym, were also associated with lower plasma GDF15 (β=-0.31, CI=-0.36 to-0.26, SE = 0.025, p=1.35×10^-35^), replicating our present findings. Given that GDF15 is the most upregulated circulating protein in human aging, these findings also align with studies of epigenetic age demonstrating that higher education level ^90,91^ and higher social support ^92^ are associated with lower biological age. GDF15 is also a marker of current and future disease risk ^89^. Future studies are required to investigate whether psychosocial factors interact with GDF15 biology to influence aging trajectories.

Our study presents several strengths including a 5-year follow-up assessment and the direct comparison of both plasma and serum cf-mtDNA with blood GDF15. However, limitations include the small group of individuals with diabetes, and that BMI and diabetes status were self-reported. Therefore, future studies are needed to replicate and extend these findings. Overall, our results establish that GDF15 and plasma cf-mtDNA are independently regulated markers of mitochondrial signaling differentially linked to aging. Based on these and other results GDF15 emerges as a particularly robust age-related biomarker of tissue stress ^93^, which can be deployed to examine the potential age-modifying influence of metabolic and psychobiological stressors.

## 4. Methods

### 4.1. Participants

Data and biospecimens were obtained from the Reference Ability Neural Network/Cognitive Reserve (RANN/CR) study ^46,94,95^. This sample included 430 community-dwelling adults (mean age = 55.0Lyears, SD = 16.7 years, range = 24–84Lyears; 54.2% women). Exclusion criteria included medical or psychiatric conditions that could affect cognition, presence of mild cognitive impairment or dementia. The Columbia University Institutional Review Board approved the Study (Protocol AAAI2752.). Participants provided informed consent before participating.

### 4.2. Procedures

A venous blood draw was taken at baseline and follow-up during the study visits. Plasma and serum were collected using EDTA and silicone-coated red-top tubes, respectively, and were extracted by centrifugation before being stored at-80°C.

Questionnaires were completed by the participants at each of the study visits. Height, weight, and type 2 diabetes status was determined from self-reports. Education levels was determined by the number of years of education. Perceived stress was measured with the perceived stress scale ^96^ and social support by the Perceived Social Support questionnaire ^97^ at the study follow-up. Physical activity was measured using Godin Exercise Questionnaire ^98^ questionnaire. Leisure activity and functional activity were measured by the Leisure Activities Questionnaire ^99^ and the Blessed Functional Activity Scale (BFAS) ^100^ respectively.

### 4.3. cf-mtDNA

cf-mtDNA in serum and plasma was quantified using the MitoQuicLy method^26^. Briefly, 10 µL of each sample was diluted into 190 µL of lysis buffer and thermolysed overnight on 96-well plates in duplicate. 8 µL of each lysate was transferred to three wells of a 384-well plate preloaded with 12 µL of a buffer containing TaqMan Fast Advanced Master Mix (Applied Biosystems, 4444557) and qPCR primers and probes targeting the mitochondrial gene ND1 and the nuclear gene B2M. qPCR was run in duplicate 384-well plates using a QuantStudio 7 Flex real-time qPCR system. For each sample, the three cycle-threshold (C_T_) measurements for each gene were averaged. If the coefficient of variance (CV) of the three measurements was greater than 0.5%, the C_T_ value that was furthest from the average was rejected and the two remaining measurements were averaged. Average C_T_ values were converted to copies of target genes per reaction by comparing them to the C_T_s of an eight-step 1:4 serial dilution series of a purified human fibroblast DNA standard (∼1.6E6 to 99 copies per reaction) which was loaded in triplicate along with samples on each qPCR plate. Values were corrected for dilution of samples into qPCR and lysis buffers to determine copies per µL sample. Finally, replicate measurements from pairs of 384-well plates were log transformed and averaged to determine a single value for each gene in each sample. Plates of samples were re-analyzed starting from lysis if the median of between-replicate CVs of log-transformed ND1 measurements was ≥ 4.0%.

### 4.4. GDF15

Plasma GDF15 concentrations were quantified using high-sensitivity ELISA kits (R&D Systems, SGD150) per manufacturer’s instructions. Samples were diluted 1:4 with assay diluent to ensure maximal count of samples within the assay’s dynamic range. Samples were run in duplicates, with each replicate on a separate plate. Absorbance was read at 450nm, and GDF15 concentrations were interpolated using a Four Parameter Logistic Curve (4PL) model in GraphPad Prism (version 9.4.1). Final concentrations from each sample was calculated by averaging the concentrations calculated across the duplicates. Samples with a C.V. greater than 15% between duplicates were re-run. Five standard curve points and three plasma reference samples were run on each plate to monitor the inter-assay C.V. and failed runs by comparing all standard curves and reference sample concentrations across all plates. R Software (version 4.2.2) was used for data pre-processing and quality control.

### 4.5. Statistical analysis

Non-parametric signed-rank Wilcoxon paired t-test was used to compare GDF15 and cf-mtDNA levels between the initial visit and the 5-year follow-up. Change in levels of plasma cf-mtDNA and GDF15 were calculated by subtracting the follow-up levels from the baseline levels, Spearman rank correlations were used to assess continuous associations. We removed one magnitude of change GDF15 value which was > 3 standard deviations away from the mean.

Non-parametric Mann-Whitney t-test was used to assess group difference. Statistical analyses were conducted using GraphPad Prism (version 9.4.1) and R Software (version 4.2.2 and 4.3.0).

## Acknowledgments

This study was supported by the NIH grants R01AG026158 and R01AG038465 to Y.S. and C.G.H., by the DRC TBAC Mini-Pilot Grants to C.T. and the Baszucki Prize in Science to M.P..

## Author’s Contribution

C.T. and M.P. conceived and supervised this research project. Y.S. and C.G. H. designed the RANN-CR study and supervised data collection. J.M., D.S., and C.C.L. performed the GDF15 and cf-mtDNA assays. C.T., J.M. and Q.H. performed statistical analyses and prepared the figures. C.T., Q.H. and M.P. drafted the manuscript. Y.S. and C.G.H. advised on manuscript and figure preparation. All authors reviewed, commented and edited the final version of the manuscript.

