## Supplemental Figures for "Blood mitochondrial health markers cf-mtDNA and GDF15 in human aging"

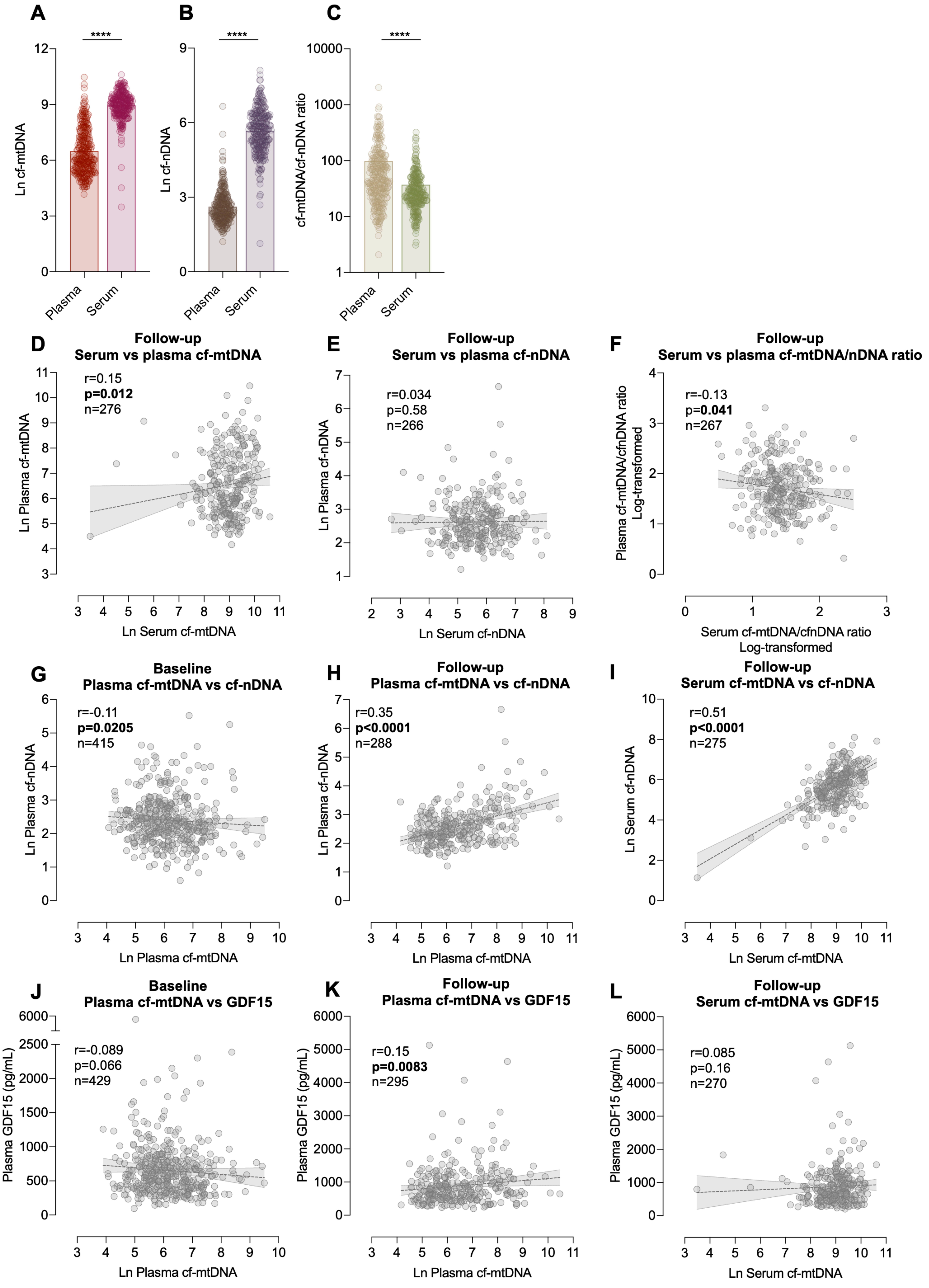


**Figure S1. Associations between serum and plasma cf-mtDNA, cf-nDNA, and GDF15 levels.** (**A-C**) Comparison between plasma and serum cf-mtDNA (**A**), cf-nDNA (**B**), and cf-mtDNA/cf-nDNA ratio (**C**) levels at the follow-up visit. (**D**) Associations between follow-up serum and plasma cf-mtDNA levels. (**E**) Same as D but for cf-nDNA levels. (**F**) Same as D but for cf-mtDNA/cf-nDNA ratios. (**G**) Associations between baseline plasma cf-mtDNA and cf-nDNA levels. (**H**) Same as **G** but for the follow-up visit. (**I**) Same as H but for serum. (**J**) Associations between baseline plasma cf-mtDNA and GDF15 levels. (**K**) Same as **J** but for follow-up visit. (**L**) Associations between follow-up serum cf-mtDNA and plasma GDF15 levels. P-value and effect sizes from (**A-C**) Mann-Whitney t-test and (**B-L**) Spearman rho correlation. *p<0.05, **p<0.01, ***p<0.001, ****p<0.0001.

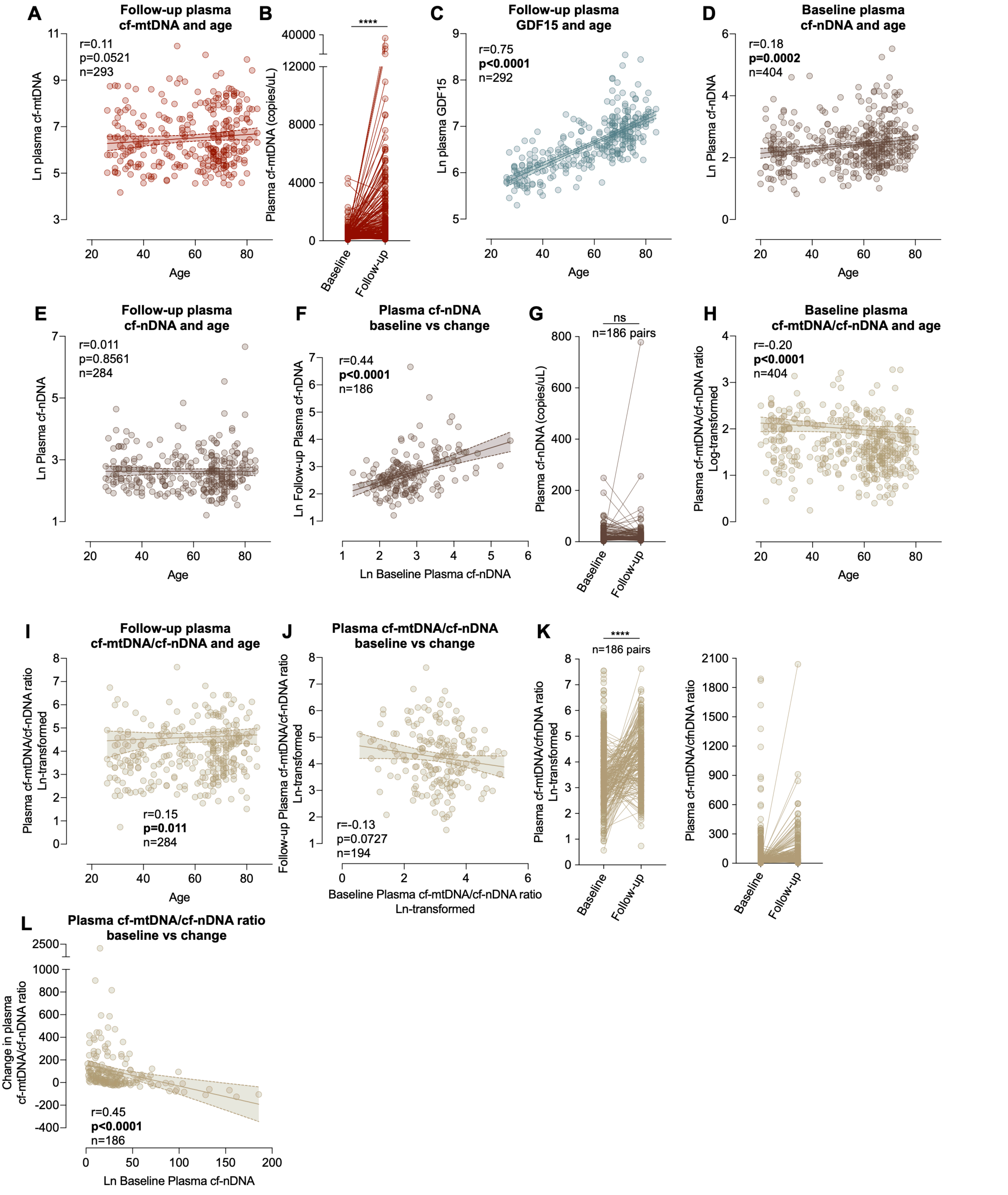


**Figure S2: cf-mtDNA, cf-nDNA, cf-mtDNA/cf-nDNA ratio associations with age and within-individual trajectories over 5 years. (A)** Scatterplot of the association between plasma cf-mtDNA levels and age at follow-up. (**B**) Change in plasma cf-mtDNA levels from baseline to follow-up in raw concentrations. **(C)** same as **A** but for plasma GDF15. (**D**) Scatterplot of the association between plasma cf-nDNA levels and age at baseline. (**E**) Same as **D** but for follow-up visit. (**F**) Scatterplot of the association between plasma cf-mtDNA levels at baseline and follow-up.(**G**) Change in cf-mtDNA levels from baseline to follow-up in natural log transformed (left) and raw concentrations (right). (**H, I, J, K**) Same as **D, E, F, G** but for cf-mtDNA/cf-nDNA ratio. (**L**)Scatterplot of the association between baseline and change in plasma cf-mtDNA/cf-nDNA levels from baseline to follow-up. P-value and effect sizes from (**B,G,K**) Wilcoxon paired t-test and (**A,C,D,E,F,H,I,J,L**) Spearman rho correlation and.*p<0.05, **p<0.01, ***p<0.001, ****p<0.0001.

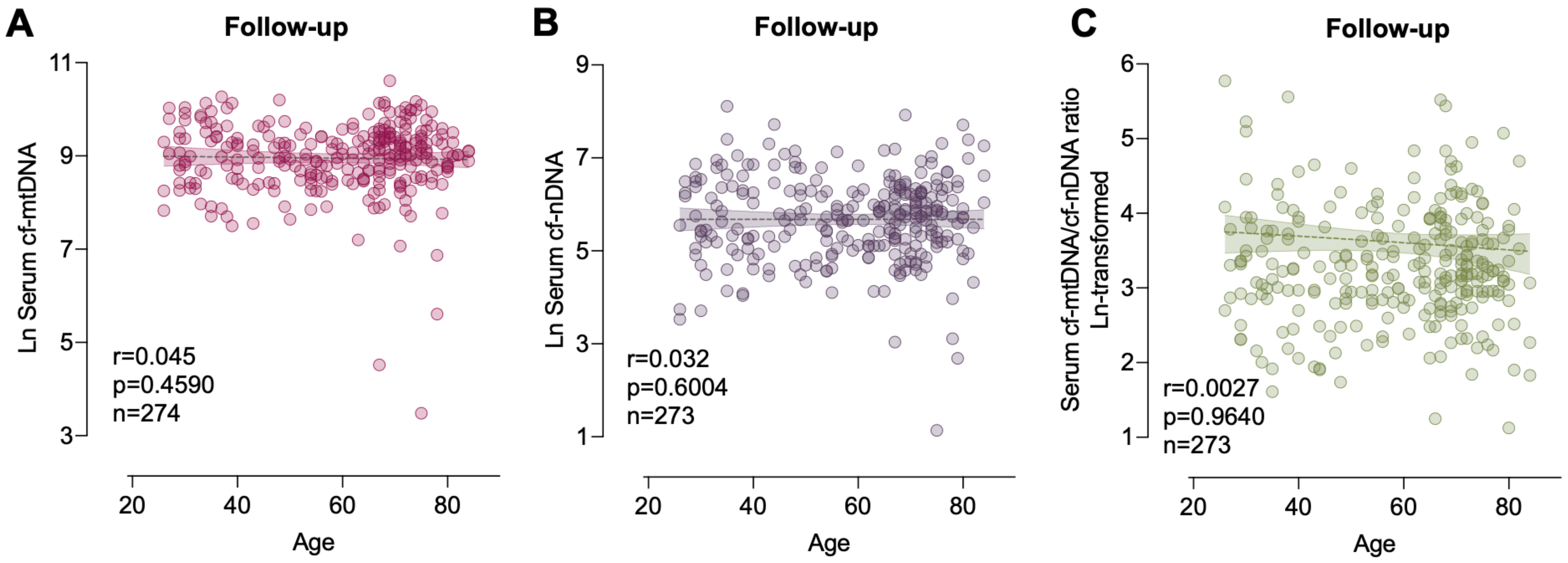


**Figure S3: Serum cf-mtDNA, cf-nDNA, cf-mtDNA/cf-nDNA ratio associations with age at the follow-up visit. (A)** Scatterplot of the association between plasma cf-mtDNA levels and age at baseline, for raw concentrations in copies/uL see Fig. S2A. (**B**) Scatterplot of the association between age and plasma cf-mtDNA levels. Same as in **B** for (**C**) plasma cf-nDNA levels, (**D**) plasma cf-mtDNA/cf-nDNA ratios, (**E**) serum cf-mtDNA levels, (**F**) serum cf-nDNA levels, (**G**) serum cf-mtDNA/cf-nDNA ratios, (**H**) plasma GDF15 levels. P-value and effect sizes from Spearman rho correlation.*p<0.05, **p<0.01, ***p<0.001, ****p<0.0001.

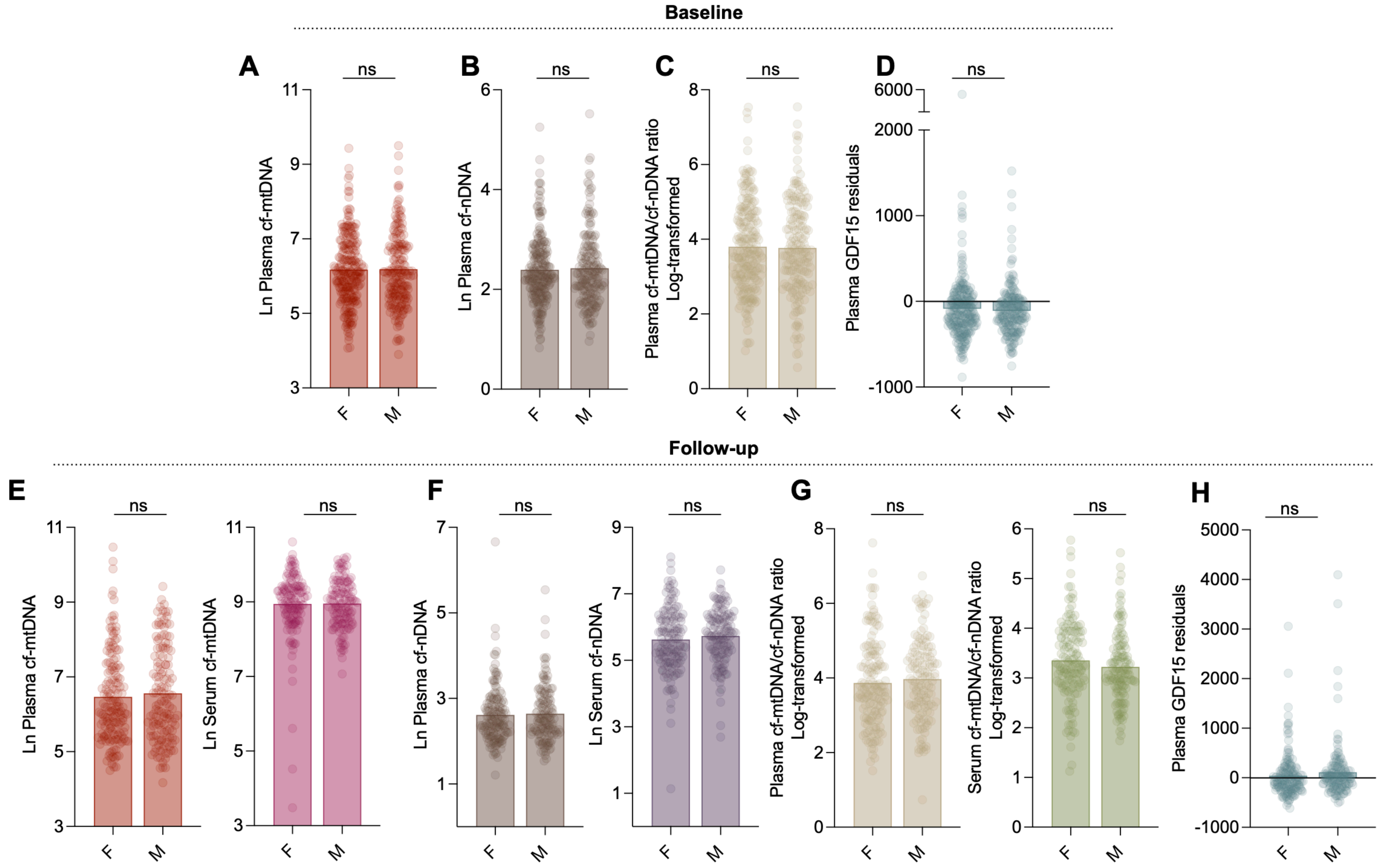


**Figure S4. Plasma and serum cf-mtDNA, cf-nDNA, cf-mtDNA/cf-nDNA ratio, and GDF15 levels by sex.** Sex difference in baseline (**A**) plasma cf-mtDNA levels, (**B**) plasma cf-nDNA levels, (**C**) plasma cf-mtDNA/cf-nDNA ratios, (**D**) plasma GDF15 levels; follow-up (**E**) plasma (left) and serum (right) cf-mtDNA levels, (**F**) plasma (left) and serum (right) cf-nDNA levels, (**G**) plasma (left) and serum (right) cf-mtDNA/cf-nDNA ratios, (**H**) plasma GDF15 levels. P-values from Mann-Whitney t-test. F = female, M = male. *p<0.05, **p<0.01, ***p<0.001, ****p<0.0001.

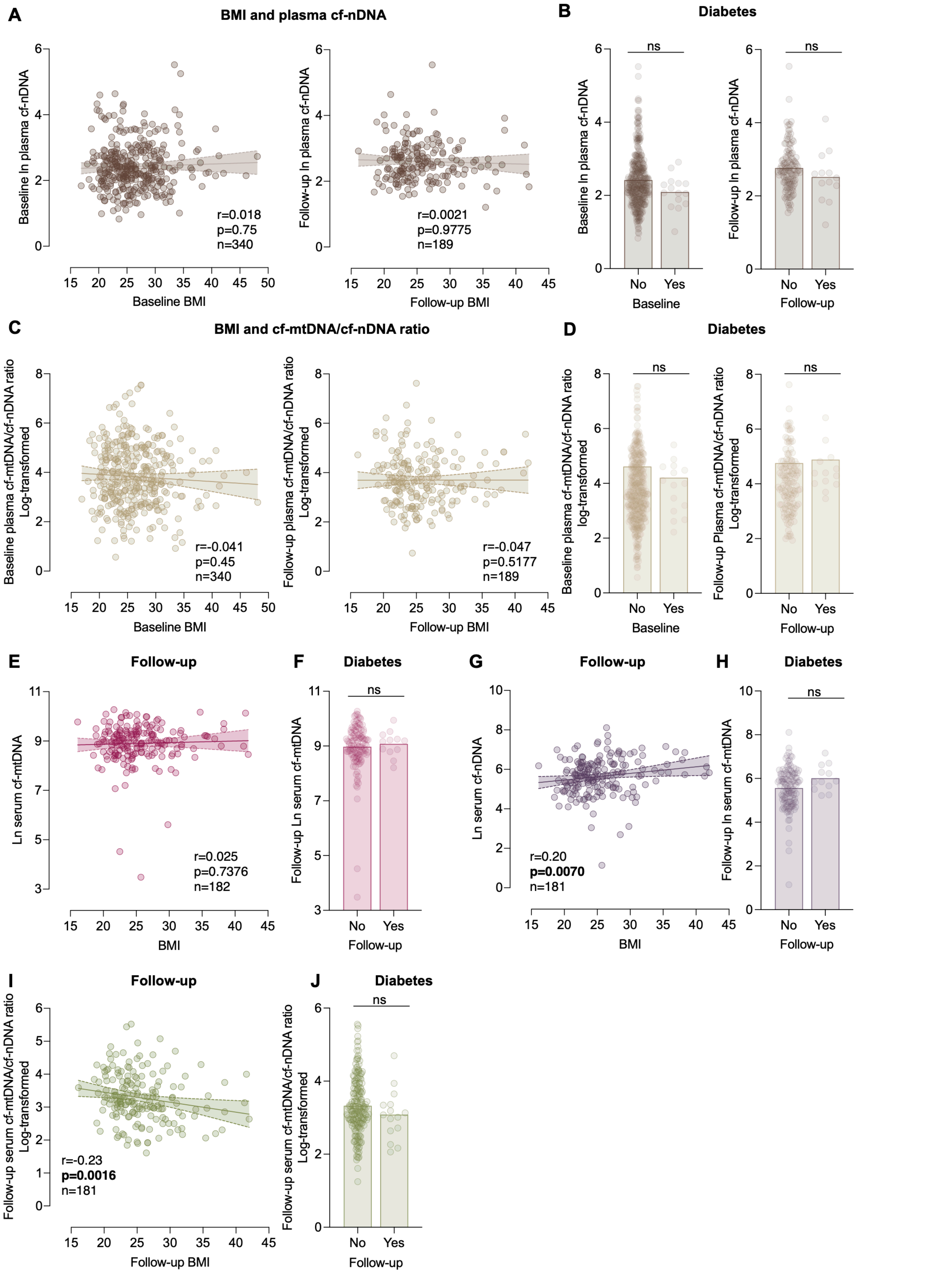


**Figure S5: Associations of cf-mtDNA levels, cf-nDNA levels, cf-mtDNA/cf-nDNA ratios with BMI and diabetes in plasma and serum**(**A**) Scatterplot of the association between plasma serum cf-nDNA levels and BMI at the follow-up visit. (**B**) Difference in plasma cf-nDNA levels by diabetic status at the baseline (left, n=389 vs 15) and follow-up (right, n=131 vs 13) visit. (**C**,**D**) Same as (**A**,**B**) but for plasma cf-mtDNA/cf-nDNA ratio, diabetic status at baseline (left, n=389 vs 15) follow-up(right, n=131 vs 13) visit. (**E**) Scatterplot of the association between serum cf-mtDNA levels and BMI at the follow-up visit. (**F**) Difference in serum cf-mtDNA levels by diabetic status at the follow-up visit, n=121 vs 12. (**G**,**H**) same as (**E**,**F**) but for serum cf-nDNA levels, diabetic status at follow-up visit n=120 vs 12. (**I**,**J**) same as (**E**,**F**) but for serum cf-mtDNA/cf-nDNA ratios, diabetic status at follow-up visit n=120 vs 12. P-value and effect sizes from (**A**,**C**,**E**, **G**, **I**) Spearman rho correlation and (**B**,**D**,**F**, **H**, **J**) Mann-Whitney t-test.*p<0.05, **p<0.01, ***p<0.001, ****p<0.0001. BL: baseline visit. F1: Follow-up visit.

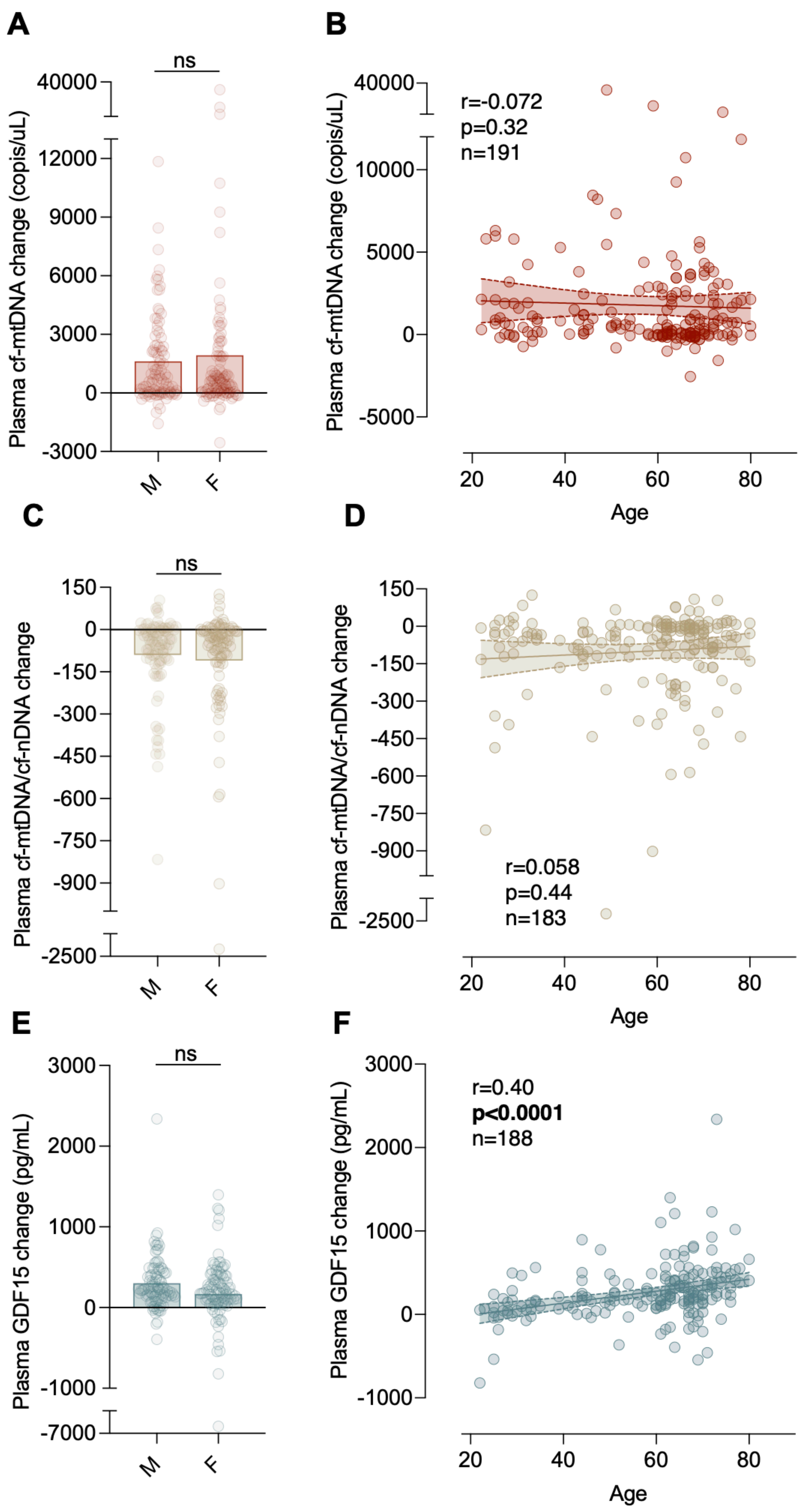


**Figure S6: Association between change in plasma cf-mtDNA, cf-mtDNA/cf-nDNA ratio, and GDF15 levels from baseline to follow-up with sex and age.** (**A**) Change in plasma cf-mtDNA levels from baseline to follow-up with by sex. Scatterplot of the association between change in plasma cf-mtDNA levels from baseline to follow-up with (**B**) age at baseline. (**C-D**) Same as in (**A-B**) but for change in plasma cf-mtDNA/cf-nDNA ratios. (**E-F**) Same as in (**A-B**) but for change in plasma GDF15 levels. P-value and effect sizes from (**A,C,E**) Mann-Whitney t-test and (**B,D,F**) Spearman rho correlation.*p<0.05, **p<0.01, ***p<0.001, ****p<0.0001.

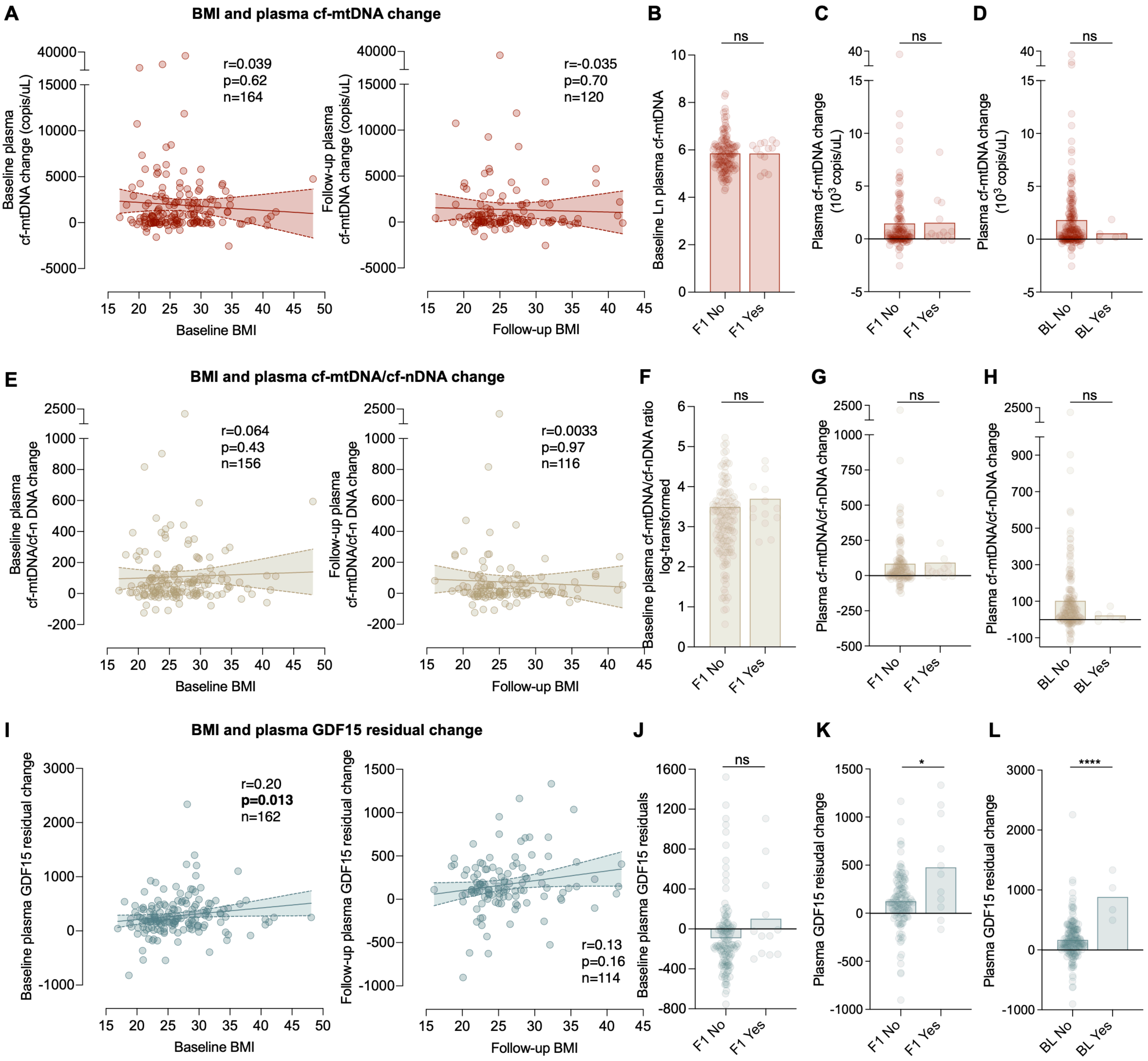


**Figure S7: Baseline and longitudinal changes in plasma cf-mtDNA, cf-mtDNA/cf-nDNA ratio, GDF15 and their relationship with BMI and diabetes status.** (**A**) Scatterplot of the association between change in plasma cf-mtDNA levels and BMI at the baseline (left) and follow-up (right) visit. (**B**) Difference in baseline plasma cf-mtDNA levels by diabetic status at the follow-up visit (n=135 vs 13). (**C**) Difference in longitudinal changes in plasma cf-mtDNA levels by diabetic status at the follow-up visit (n=135 vs 13). (**D**) Association between diabetic status at baseline (n=186 vs 5) and the longitudinal changes in plasma cf-mtDNA levels. (**E-H**) Same as in (**A-D**) but for change in plasma cf-mtDNA/cf-nDNA ratios. n=134 vs 13 in D, n=130 vs 13 in E, n=178 vs 5 in F. (**I-L**) Same as in (**A-D**) but for change in plasma GDF15 residuals. n=132 vs 12 in G, n=131 vs 11 in H, m=184 vs 4 in I. P-value and effect sizes from (**A,E,I**) Spearman rho correlation and (**B,C,D,F,G,H,J,L**) Mann-Whitney t-test and.*p<0.05, **p<0.01, ***p<0.001, ****p<0.0001. BL: baseline visit. F1: Follow-up visit.

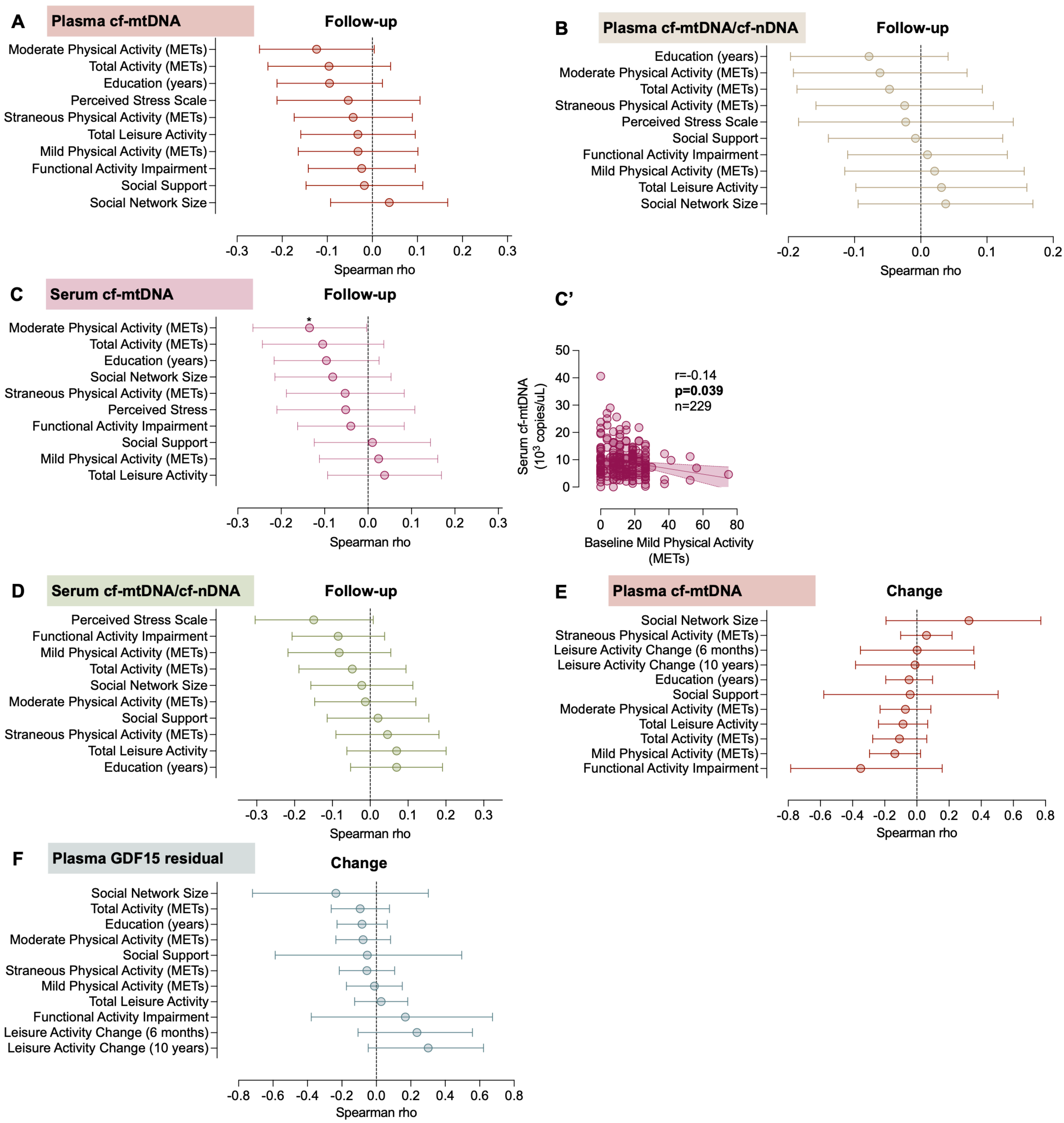


**Figure S8: Associations between plasma and serum cf-mtDNA, cf-mtDNA/cf-nDNA ratio, GDF15 and behavioral and psychosocial factors at baseline and follow-up**

(**A**) Associations between follow-up plasma cf-mtDNA levels with behavioral and psychosocial factors ranked from most negative to most positive. (**B**) Same as in **A** but for follow-up plasma cf-mtDNA/cf-nDNA ratios. (**C**) Same as in A but for follow-up serum cf-mtDNA levels. See (**C'**) for statistically significant correlations with Mild Physical Activity (METs). (**D**) Same as in **A** but for serum cf-mtDNA/cf-nDNA ratios. (**E**) Associations between change in cf-mtDNA levels with bahavioral and psychosocial factors at baseline ranked from most negative to most positive. (**F**) Same as in **E** but for change in plasma GDF15 residuals. P-value and effect sizes from Spearman rho correlation. *p<0.05, **p<0.01, ***p<0.001, ****p<0.0001.
